# Antibiotic Resistance in *Bacillus*-based Biopesticide Products

**DOI:** 10.1101/2021.03.15.435560

**Authors:** Mo Kaze, Lauren Brooks, Mark Sistrom

**Author notes:** Corresponding author Mo Kaze, UC Merced, 5200 N Lake Rd, Merced, CA 95343. CLASSIFICATION: Applied and Environmental Science, Microbiology.

## Abstract

The crisis of antibiotic resistant bacterial infections is one of the most pressing public health issues. Common agricultural practices have been implicated in the generation of antibiotic resistant bacteria. Biopesticides, live bacteria used for pest control, are non-pathogenic and considered safe for consumption. Application of bacteria-based pesticides to crops in high concentrations raises the possibility of unintentional contributions to the movement and generation of antibiotic resistance genes in the environment. However, the presence of clinically relevant antibiotic resistance genes and their resistance phenotypes are currently unknown. Here we use a combination of multiple bioinformatic and microbiological techniques to define resistomes of widely used biopesticides and determine how the presence of suspected antibiotic resistance genes translates to observable resistance phenotypes in several biopesticide products. Our results demonstrate that biopesticide products are reservoirs of clinically relevant antibiotic resistance genes and bear resistance to multiple drug classes.

**Importance:** This is the first study to specifically address antibiotic resistance in widely distributed bacterial strains used as commercial biopesticides. Safety assessments of commercial live bacterial biopesticide products do not include antibiotic resistance phenotype identification. We identify antibiotic resistance genes in all live bacterial strains examined, and resistant phenotypes in all strains tested for antibiotic susceptibility. This work demonstrates that biopesticides potentially play a critical role as reservoirs and vectors of antibiotic resistance in the broader environmental resistome that is to date, unstudied.

## Introduction

The increasing prevalence of antibiotic resistant bacterial infections is one of the most pressing public health crises of the current era. Without significant efforts to curb antibiotic resistant infections, 10 million human deaths per annum are estimated to occur by 2050 along with severe impacts to animal husbandry and subsequent food production.(1–2) A large body of research has investigated the spread of antibiotic resistance via common agricultural practices, implicating many in the generation of antibiotic resistant bacteria.(1–6) The widespread use and presence of antibiotic compounds in the broader environment, and the ubiquitous presence of genes encoding resistance to them, play a critical role in the evolutionary mechanisms affecting antibiotic resistance. Clinically relevant and nonclinical bacterial species residing in microorganismal communities horizontally transfer resistance genes. These exchanges subsequently influence the prevalence and patterns of antibiotic resistant infections.(7,8)

The use of microorganisms in agriculture for pest control, frost prevention, and rhizosphere enhancements has steadily increased over the last 20 years.(9,10) Considered safe for consumption, non-toxic, non-pathogenic and highly effective,(11–14) microbial products offer a welcome alternative to chemically synthesized pesticides known to cause damage to human health and the environment.(15, 16) Microbes used as biopesticides are classified “Generally recognized as safe” (GRAS) by the US FDA(17) as they do not pose a threat to human health. However, these microorganisms have the potential to contribute to the pervasiveness of antibiotic resistance through genes encoded in bacterial genomes and mobile genetic elements. It is crucial to identify clinically relevant antibiotic resistance genes present in live bacterial biopesticides used in large scale applications to prevent unintentional contributions to the spread of antibiotic resistance genes and the expansion of antibiotic resistance gene reservoirs. There is an urgent need to understand the role biopesticides play in the transmission of antibiotic resistance genes and their roles as potential vectors.

*Bacillus*-based biopesticides are increasingly popular. *Bacillus thuringiensis* (Bt) is the most widely used biopesticide in industrial agriculture. Aerial Bt spraying has replaced aerial DDT, a known environmental toxin, for control of moths, blackflies, mosquitoes, and many other pests in forestry, agriculture, and urban areas.(18,19) Out of commercial *Bacillus*-based biopesticides, Bt is considered the safest, and has been in use globally for more than 80 years.(20) Bt is a Gram positive, aerobic, soil-dwelling bacteria characterized by the presence of plasmids containing *cry* and *cyt* genes.(21) These two toxin genes and their variations confer unique insecticidal properties. Bt is genetically plastic and has special capability regarding plasmid acceptance and maintenance. This biopesticide species has previously been shown to host as many as seventeen plasmids.(22)

This study represents the first effort to assess the potential role of live, commercial bacterial biopesticides as reservoirs of antibiotic resistance genes, and to connect antibiotic resistance phenotypes to resistance genotypes. We analyze four commercially available *Bacillus*-based biopesticide strains: two Bt kurstaki products, *B*. *amyloliquefaciens* D747, and *B. subtilis* QST 713, using a combination of bioinformatics and antibiotic susceptibility testing. We classify all antibiotic resistance genes in these *Bacillus*-based biopesticide products by comparing whole-genome sequenced products against the Comprehensive Antibiotic Resistance Database (CARD) and annotating sequenced genomes.(23) This work demonstrates that currently used commercial *Bacillus*-based biopesticides contain clinically relevant antibiotic resistance genes and bear resistance to multiple drug classes. These findings raise concern regarding potential vectors of unintended transmission of antibiotic resistance as they are introduced to the environment in large quantities.

## Methods

A list of all *Bacillus*-based biopesticide species approved for use in the US was collated from databases published by the California Department of Pesticide Registration and United States Environmental Protection Agency databases.(24) In order to narrow down antibiotic selection, publicly available complete, whole reference genomes matching strain information were queried against the Comprehensive Antibiotic Resistance Database v3.0 (downloaded November 2019) using Blastn 2.10.0(25) with default settings, except for a 97% cut off for query coverage and 97% percent for percent identity match. Reference genome annotations were manually reviewed for antibiotic resistance genes and proteins. Drug classes that were not present in the reference genomes were not included for phenotype testing.

Four commercial *Bacillus*-based biopesticide products, Bt subspecies kurstaki strain SA12, *B. subtilis* strain QST 713 and *B. amyloliquefaciens* strain D747, were purchased and named: Bt-kurstaki-1, Bt-kurstaki-2, *B. subtilis*, and *B. amyloliquefaciens*. The two Bt products were the same strain, but from different companies and contained different suspension materials. Commercial biopesticide products were cultured using standard methods and McFarland turbidity as specified in the American Society of Microbiology Kirby-Bauer Disk Diffusion Susceptibility Test Protocol(26) and assayed with Oxoid (Thermo Fisher, USA) antimicrobial susceptibility disks for clindamycin (2 μg), doxycycline (30 μg), linezolid (30 μg), sulfamethoxazole/trimethoprim (25 μg), and vancomycin (30 μg). Minimum inhibitory concentration (MIC) was determined using the standard Clinical and Laboratory Standards Institute guidelines(27) on replicates using Liofilchem (Liofilchem Inc. MA) antibiotic minimum inhibitory concentration test strips and aerobe incubation protocols for cephazolin (0.016 – 256 μg/mL), clindamycin (0.016 – 256 μg/mL), ceftazidime (0.016 – 256 μg/mL), quinupristin/dalfopristin (0.002 – 32 μg/mL), ertapenem (0.002 – 32 μg/mL), imipenem (0.002 – 32 μg/mL), erythromycin (0.016 – 256 μg/mL), and tetracycline (0.016 – 256 μg/mL) antibiotics. MIC breakpoints were obtained from EUCAST.(28)

DNA from biopesticide products was extracted directly and from Luria-Bertani cultures using Qiagen’s DNeasy DNA Extraction kit (Qiagen NV, Germany) with a modified protocol. After performing the protocol’s first step, samples were incubated at 90°C for 10-15 minutes in order to account for Gram positive cell wall structure. DNA concentration was quantified using a Qubit fluorometer (Thermo Fisher Scientific, MA) and quality and purity quantified using an Eppendorf Biospec (Eppendorf, Germany). Libraries were generated using Illumina NextTera Flex kit with IDT set A Dual Indexes. DNA was sequenced on an Illumina MiSeq platform (Illumina Inc, CA). Sequences were checked for quality using FastQC v0.11.8(29) and trimmed using Trimmomatic v0.36. (30) Genomes were assembled using SPAdes v3.11.1.2(31) and assessed for quality using Quast v5.0.0.(32) Genomes were annotated with RAST v4.0.2(33) and assemblies were queried against CARD and manually curated for antibiotic resistance annotations and verified against UniProt. (34) Annotated and CARD-identified antibiotic resistance genes were quantified in R Studio v 1.4.1103.(35)

## Results

### Antibiotic Susceptibility

Each biopesticide product demonstrated antibiotic resistance phenotypes to the clinically relevant antibiotics tested. (**Table 1**) The MIC range for replicates for clindamycin was 0.064 μg/mL to 0.19 μg/mL and erythromycin was 0.125 μg/mL to 0.19 μg/mL. Ertapenem resistance for all products ranged from 0.125 μg/mL to 0.19 μg/mL. Imipenem resistance was observed in Bt-kurstaki-1, Bt-kurstaki-2 and *B. amyloliquefaciens* products. Bt-kurstaki-1 was interpreted as resistant with all replicates measuring 1.5 μg/mL. Bt-kurstaki-2 was resistant in 20% of replicates (sd. 1.6) and 80% were susceptible to imipenem and ranged from 0.094 μg/mL to 4.0 μg/mL. *B. amyloliquefaciens* was susceptible to imipenem in 80% of replicates and ranged from 0.032 μg/mL to 0.75 μg/mL (sd. 0.3). Resistance to quinpristin/dalfopristin was observed in *B. amyloliquefaciens* measuring 3.0 to 4.0 μg/mL (sd. 0.3) and B. subtilis measuring 3.0 μg/mL. Both of these biopesticide products were also resistant to tetracycline. Bt-kurstaki-1 and Bt-kurstaki-2 had complete resistance to both cephalosporins tested, ceftazidime and cefazolin, with MICs of 256 μg/mL. Disk diffusion assays showed resistance to two of the five antibiotics tested. (**Table 2**) Clindamycin resistance was observed in 25% of Bt-kurstaki-1 (n = 12, 0 mm). Sulfamethoxazole/trimethoprim resistance was observed in 33% of Bt-kurstaki-2 replicates (n = 9, 10 mm)*. B. subtilis* and *B. amyloliquefaciens* were susceptible to all five antibiotics. All four biopesticide products tested were susceptible to doxycycline, linezolid, and vancomycin. Resistance to five total drug classes, across the four biopesticide products, was observed: cephalosporins, lincosamides, streptogramins, sulfonamides, and tetracyclines. **(Fig. S1, Fig.S2)**

### Molecular Characterization

Genotypes for each biopesticide product contained multiple antibiotic resistance genes for the five drug classes of the observed resistance phenotypes. Both methods used to identify antibiotic resistance genes, a curated database of resistance genes and genome annotation, identified antibiotic resistance genes associated with the nine tested drug classes. For the five drug classes represented by the resistance phenotypes, CARD identified twelve genes and genome annotation identified seven. Of the fifty-two total antibiotic resistance genes identified by CARD and the forty-seven identified by genome annotation, less than half resulted in an expressed resistance phenotype (44% and 47% respectively).

CARD identified twenty-two antibiotic resistance genes associated with the antibiotics tested for susceptibility. (**Fig. 1a**) Bt-kurstaki-1 had eighteen (82% of the total genes for the nine tested drug classes) genes for resistance to eight drug classes: carbapenems (Bla2, MexB, MexY), cephalosporins (BcI, BcII, lsaB, MexB, MexD, MexY), glycopeptides (vanRM), lincosamides (lsaB), macrolides (lsB, mdtF, MexB, MexD, MexY), oxazolidinones (lsaB), sulfonamides (MexB, sul1), and tetracyclines (acrA, lsaB, MexB, MexD, MexY, MuxB, MuxC, oqxA, oqxB, smeE, tet(L)). Bt-kurstaki-2 had six genes for seven of the drug classes tested: carbapenems (Bla2), cephalosporins (BcI, BcII, lsaB), glycopeptides (vamRM), lincosamides (lsaB), macrolides (lsaB), oxazolidinones (lsaB), and tetracyclines (acrA, lsaB). *B. amyloliquefaciens* and *B. subtilis* products contained the same five genes associated with six of the tested drug classes: cephalosporins (hns), lincosamides (cfr(B), clbA, clcD), macrolides (hns), oxazolidinones (cfr(B), clbA, clcD), streptogramins (cfr(B), clbA, clcD), and tetracyclines (hns, tet(L)).

Gene annotation of sequenced biopesticide genomes identified twenty-two genes associated with the antibiotics tested for susceptibility. Bt-kurstaki-1 and *B. subtilis* contained all twenty-two genes for eight drug classes. Bt-kurstaki-2 contained fourteen (67%) genes which were associated with six drug classes: cephalosporins (AcrE, CmeABC, MarA, MarB, MarR), carbapenems (MarA, MarB, TolC), lincosamides (ErmA, ErmB), macrolides (CmeABC, ErmA, ErmB), streptogramins (ErmA, ErmB), and tetracyclines (AcrA, MarA, MarB, MarR, mdfA, tetR, TolC). *B. amyloliquefaciens* had nineteen (86%) genes associated with eight of the tested drug classes: carbapenems (MarA, MarB, TolC), cephalosporins (AcrE, CmeABC, MarA, MarB, MarR), erythromycins (mdlB), glycopeptides (vanW), lincosamides (ErmA, ErmB), macrolides (CmeABC, ErmA, ErmB, MacA, MacB, TolC), streptogramins (ErmA, ErmB), and tetracyclines (AcrA, MarA, MarB, MarR, MdfA, tetR, TolC, YkkC, YkkD). The complete list of identified antibiotic resistance genes are summarized in **Table 3**.

**Table 3.**
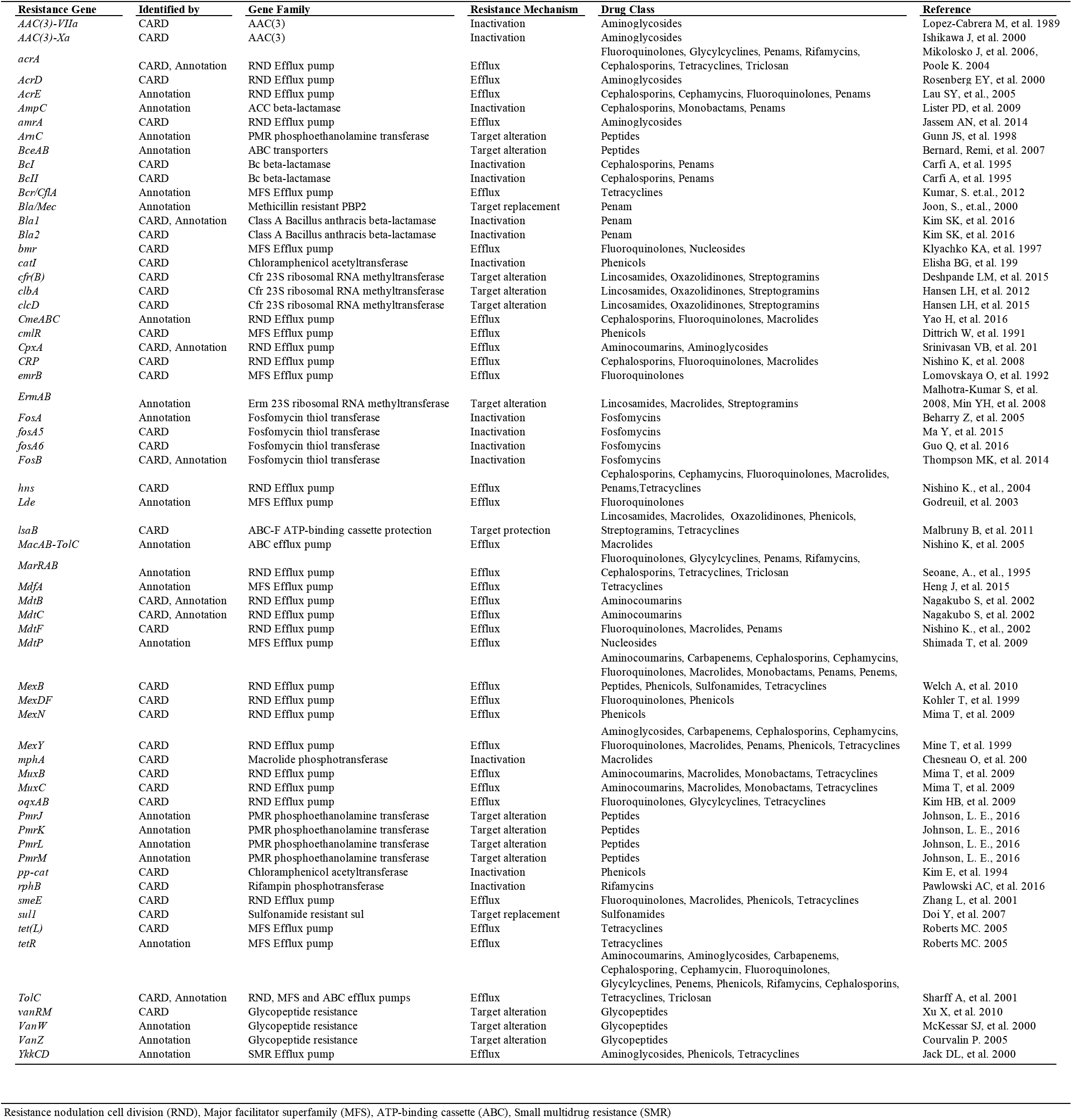
Antibiotic Resistance Genes Summary. Resistance nodulation cell division (RND), Major facilitator superfamily (MFS), ATP-binding cassette (ABC), Small multidrug resistance (SMR)

The two methods used to determine genotypes provided different results and neither method accounted for differing genotypes relating to drug class phenotype. Annotation did not identify genes for oxazolidinone and sulfonamide resistance. Seven resistance genes were identified by both methods: AcrA, Bla1, CpxA, fosB, MdtB, MdtC, and TolC. Annotation identified the majority of the genes associated with the antibiotic phenotypes observed. CARD identified twenty-one drug classes total and genome annotation identified fourteen drug classes. Comparing the two products with identical strains, eighteen genes were identified by CARD for Bt-kurstaki-1 and six in Bt-kurstaki-2, whereas annotation identified twenty-two genes in Bt-kurstaki-1 and fourteen in Bt-kurstaki-2. The quantification of total genes identified by both methods showed little variation between the two. (**Fig. S3**)

## Discussion

Agricultural practices currently implicated in the antibiotic resistance crisis do not currently encompass all processes contributing to the spread and maintenance of resistant bacteria in the environment. Biopesticides are disseminated globally in large quantities but have yet to be looked into as reservoirs of antibiotic resistance genes resulting in a lack of data regarding biopesticide-specific strains’ resistance phenotypes and accounting of their resistance genotypes. Assessing antibiotic resistance phenotypes or genotypes has not historically been included when testing the safety of biopesticide use or included in antibiotic resistance surveillance. Biopesticide products may act as latent carriers and as potential vectors of resistance to human pathogens which may not be determined by susceptibility testing of biopesticide products. This study spotlights antibiotic resistance phenotypes and genotypes in *Bacillus*-based biopesticides and signals the need for investigation of this agricultural practice acting as reservoirs of antibiotic resistance along the food chain.

*Bacillus*-based biopesticide strains harboring a variety of antibiotic resistance genes and expressing resistance to first-generation antibiotics, such as narrow spectrum beta-lactamases, is expected. However, resistance phenotypes and genotypes associated with later generation, clinically important antibiotics is cause for serious concern. The addition of large amounts of live bacteria for pest control increases the likelihood of horizontal gene exchange between pathogenic bacteria and biopesticides bearing resistance genotypes. All assayed biopesticide products demonstrated resistance phenotypes to two clinically important antibiotics. We observed resistance to five drug classes, all designated critically important by the World Health Organization: cephalosporins, carbapenems, lincosamides, streptogramins, and tetracyclines. (36) Genotypes contain genes capable of conferring resistance to additional clinically relevant antibiotics and biocides. Both *B. thuringiensis* products demonstrated complete resistance to ceftazidime, a third-generation cephalosporin. Resistance to third-generation broad spectrum cephalosporins are of special concern, the WHO has categorized this drug class as “highest priority critically important antimicrobials.”(26) Resistance to imipenem, a broad spectrum beta lactamase usually reserved for multi-drug resistant infections,(37) was found in one *B. thuringiensis* product, and in some replicates of the other *B.* thuringiensis product, as well as in *B. amyloliquefaciens* replicates.

There are previous examples of studies identifying antibiotic resistance phenotypes and genotypes in additional biopesticide genera. (38) Patel et al. identified vancomycin resistance clusters in biopesticide *Paenibacillus popillae.* (39) *Burkholderia ambifaria*, while no longer approved for biopesticide use in the United States, (40) contains genes required for resistance-nodulation-cell division (RND) efflux pumps.(40) We identified accessory genes associated with vancomycin resistance in the assayed biopesticide products, vanRM (CARD) vanW and vanZ (annotation). While the role of these genes is not currently understood, (39,42) the lack of other essential components likely explains why vancomycin resistance was not observed in any product. Luna et. al. tested six *Bacillus* species, both clinical and environmental, for antibiotic sensitivity and observed susceptibility in 100% of *B. thuringiensis* replicates to erythromycin, and vancomycin and 95% of replicates were susceptible to clindamycin and sulfamethoxazole/trimethoprim. (43) These results are similar to our resistance observations for both antibiotics tested against two *B. thuringiensis* kurstaki products. Turnbull et al. tested clinical and environmental isolates of *B. thuringiensis* and identified resistance phenotypes with MICs; 100% of isolates were resistant to cefotaxime, and 80% of isolates were resistant to tetracycline; with all susceptible to erythromycin and vancomycin. (44) Both *B. thuringiensis* kurstaki products we tested demonstrated complete resistance to ceftazidime, a third-generation cephalosporin. Turnbull et. al. isolates were resistant to 3rd generation cephalosporin cefotaxime. (41) While this study was not testing biopesticide specific strains, this report is also consistent with our findings.

Resistance interpretations were determined by comparing results to EUCAST references (28) However, breakpoints for each species of *Bacillus*-based biopesticide are not available for the majority of clinical antibiotics. When reference values were unavailable interpretations were determined by comparisons to reference values for related pathogenic *Bacillus*-species and taking into account the strength of the antibiotic dose. Defining resistance phenotypes and genotypes can readily be expanded to include more biopesticide products and additional clinically relevant antibiotics. Genes identified by both methods point to multiple drug classes that require further investigation: cpxA, mtdB, mdtC (aminocourmarins), Bla1 (penams), fosB (fosfomycins), and arcA, TolC (triclosan). Characterizing the genomes of these products offers an opportunity to define breakpoints for non-pathogenic species and test for additional antibiotic resistance.

Despite vociferous support for *B. thuringiensis* as the “safest … microbial insecticide available to humanity,” (45) antibiotic resistance phenotypes for critically important drug classes and the potential to share resistance conferring genes via horizontal gene transfer have not been included in any safety assessment. We observed individual mechanism genes for incomplete RND efflux pumps, e.g., CARD identified only smeE, a member of the complex for a multidrug RND efflux pump (46) as well as muxB and muxC which require genes muxA and OpmB to function. (47) TolC was identified by both methods, in all four biopesticide genomes. This gene is an essential component of multiple antibiotic resistance gene families: ATP-binding cassette, major facilitator superfamily, and RND antibiotic efflux pumps. (48) While inactive on their own, genetic exchange between strains may generate additional phenotypes as individual genes combine to form functional resistance mechanisms before application. Plasmids have been found to have very large host ranges and genetically plastic *Bacillus* species are able to host many plasmids. (49) Multiple genes identified in the biopesticides tested were initially found in mobile genetic elements: clbA is a cfr gene found in *B. amyloliquefaciens* subsp. plantarum plasmids (50)and sul1, confers sulfonamide resistance, is associated with integrons.(4, 51) It is likely that live bacterial biopesticides come into contact with, and exchange with, other genetically plastic, agriculture associated, antibiotic resistant pathogens such as *Klebsiella pneumoniae* or *Escherichia coli*.(4, 52) TolC has been associated with resistance to fifteen drug classes and has been identified in resistant *K. pneumoniae* and *E. coli*.(53, 54) Including additional methods for genotype characterization, such as use of additional curated databases,(55) may assist in predicting potential vector activity before combining biopesticides. As biopesticides are commonly used in multi-strain consortia it will be important to experimentally determine the capability for exchange between the biopesticides themselves and exchange with agriculturally associated pathogens.

Production of *B. thuringiensis* strains for pest control without creating resistance to broad spectrum antibiotics was proposed as early as 1997, (56) indicating prior concern regarding application of biopesticides acting as antibiotic resistance gene reservoirs. Despite this initial concern, the assertions of a large number of studies regarding the safety of live bacterial pesticides, specifically *B. thuringiensis* strains, mention antibiotic resistance as a factor in their evaluations.(11–14, 19, 20) Anthropogenic input of resistant bacteria in proximity to the largest users of antibiotic compounds has the potential to be a major driver in the increased prevalence of clinically relevant antibiotic resistance genes. To date, this agricultural activity, remains largely uninvestigated. Globally, live bacterial biopesticide products play an essential role in replacing toxic, chemically synthesized pesticides and offer an inexpensive and effective method of pest control. However, our results show the potential of biopesticides as a large reservoir for the generation and dissemination of antibiotic resistance genes on a global scale. While biopesticides have a role in providing safe food production, we have demonstrated that they also may play a role in the antibiotic resistance crisis. We anticipate the role of biopesticides acting reservoirs and vectors of clinically relevant antibiotic resistance genes to become part of the continued research regarding agricultural practices and the generation and spread of antibiotic resistance in the environment. Globally, live bacterial biopesticide products play an important role in replacing toxic, chemically synthesized pesticides and offer an inexpensive and effective method of pest control. However, our results show the potential of biopesticides as a large reservoir for the generation and dissemination of antibiotic resistance genes on a global scale. While biopesticides play an important role in providing safe food production, we have demonstrated that they also maybe play an important role in the antibiotic resistance crisis.

---

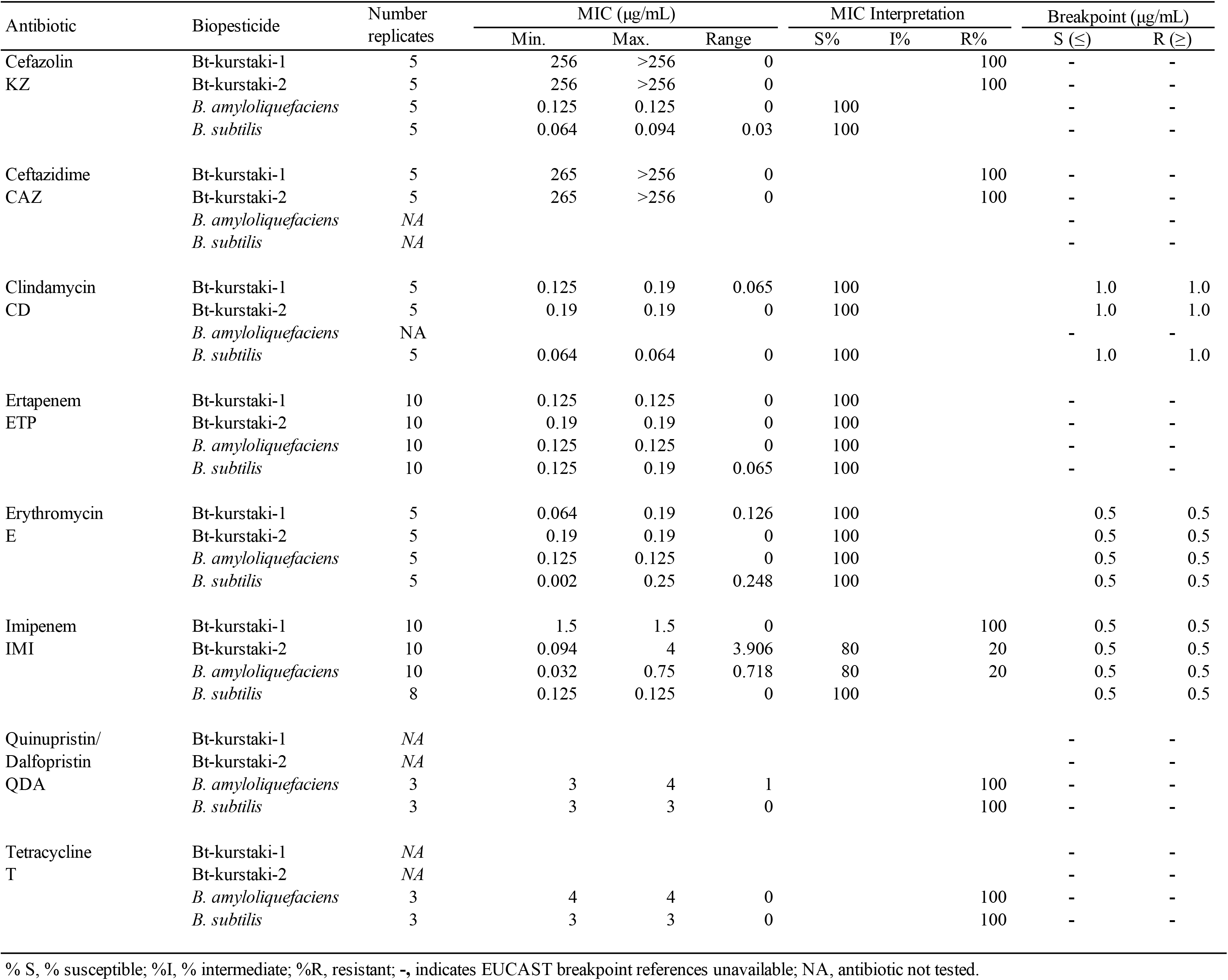

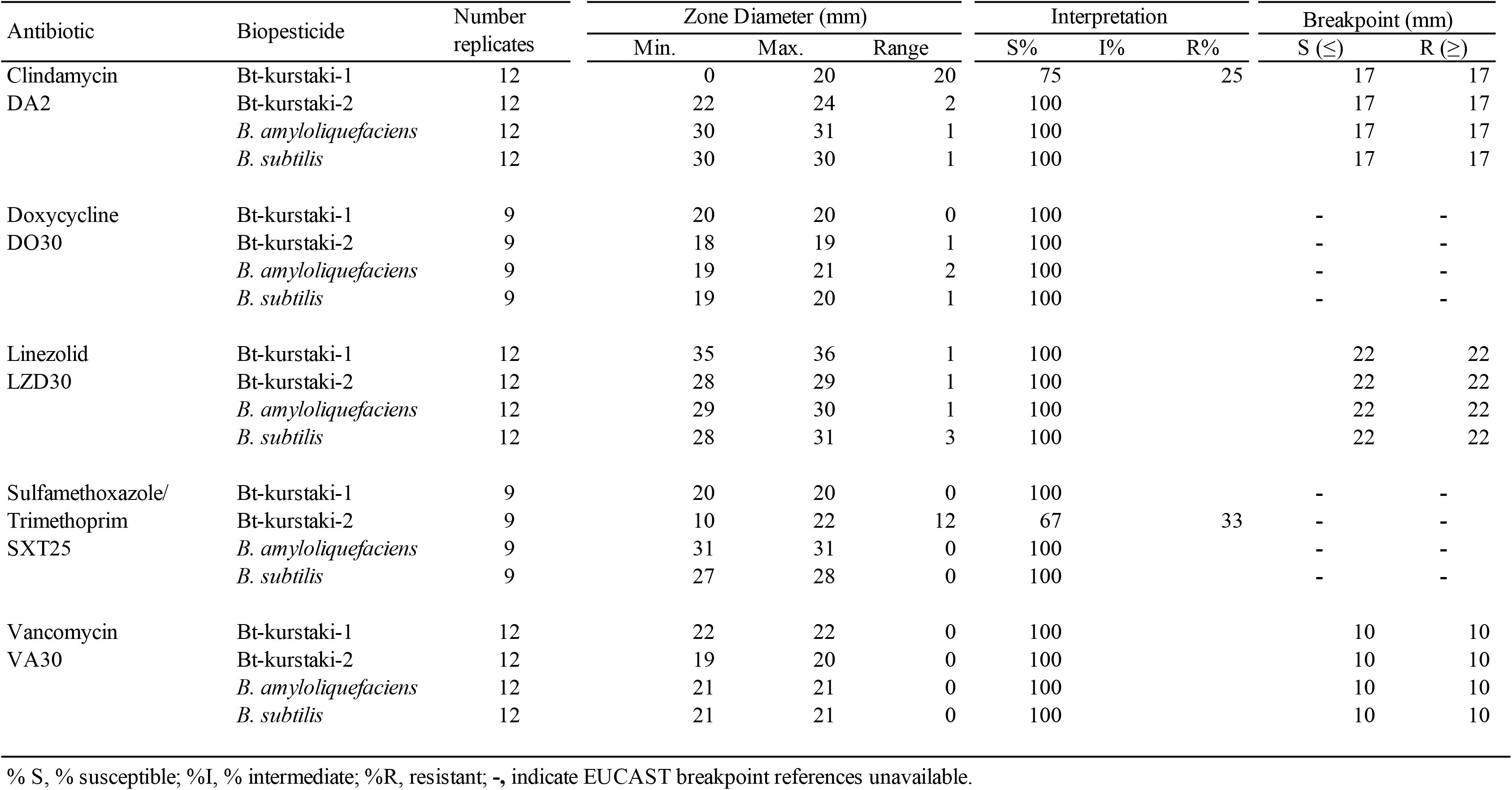

**Figure 2.**
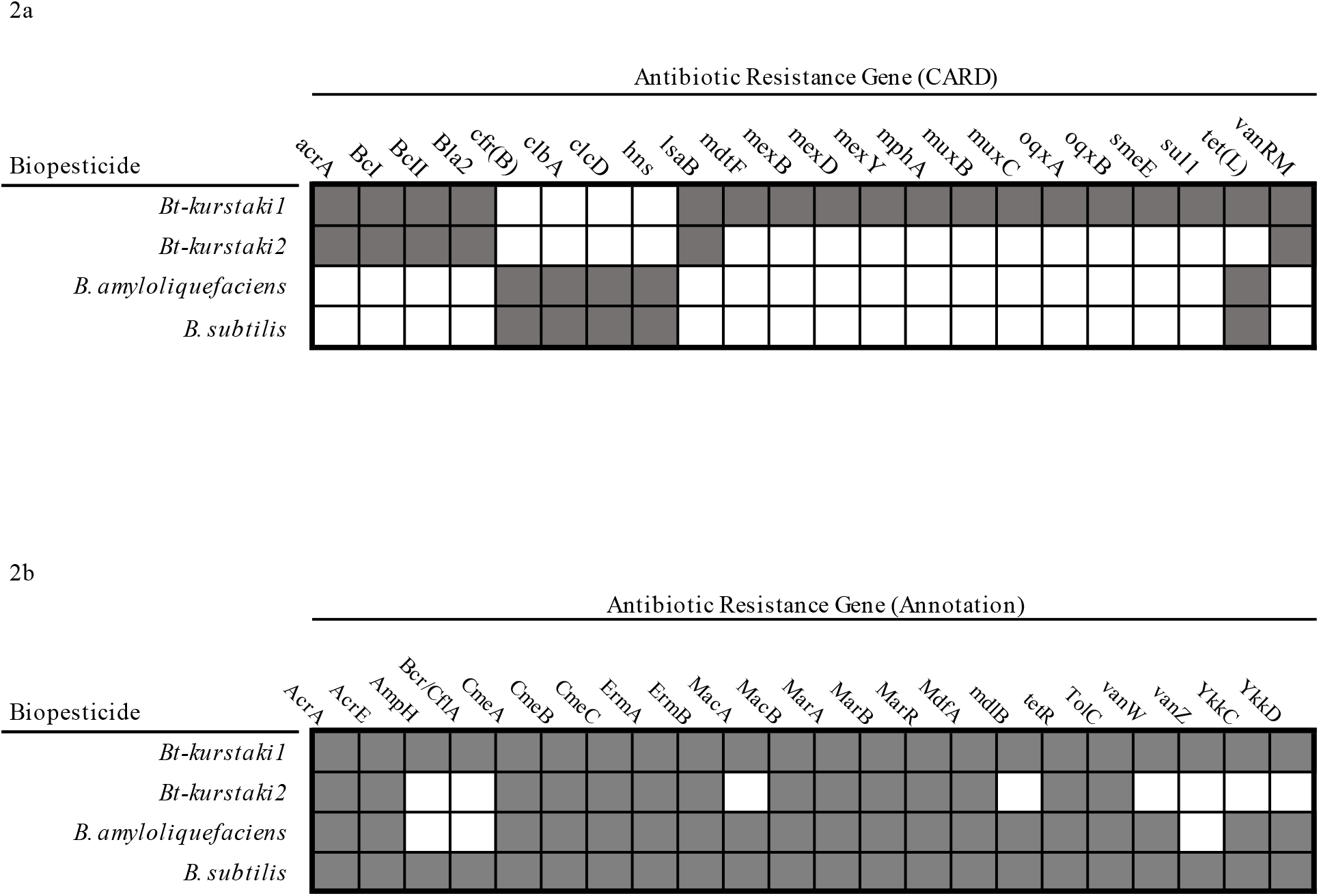
Molecular Characterization of Bacillus-based Biopesticides. Molecular characterization of commercial biopesticides showing the presence of antibiotic resistance genes associated with each drug class resistance phenotypes. **a**. CARD-identified ARG for each commercial biopesticide product. **b**. Annotation-identified ARG for each commercial biopesticide. Dark squares indicate gene presence.

**Figure S1.**
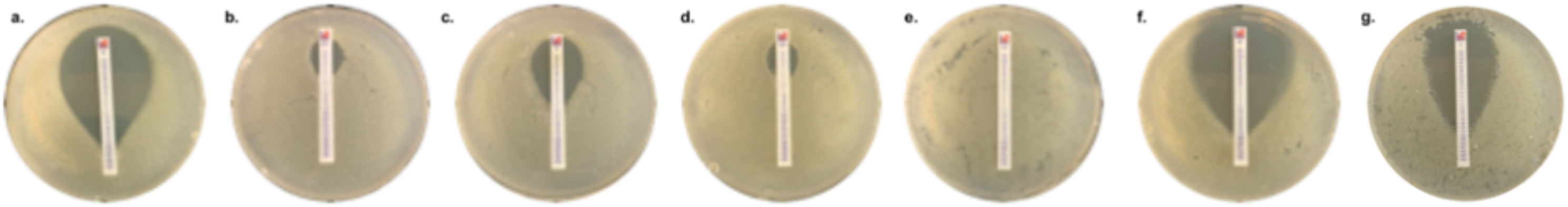
Selection of MIC results on Mueller Hinton agar for the *Bacillus*-based biopesticide products. **a.** *Bacillus subtilis* and cephazolin **b.** *Bacillus subtilis* and quinupristin/dalfpopristin. **c.** *Bacillus subtilis* and tetracycline. **d.** *Bacillus amyloliquefaciens* and quinupristin/dalfpopristin. **e.** Bt-kurstaki-2 and ceftazidime. **f.** Bt-kurstaki-2 and cephazolin.**g.** Bt-kurstaki-1 and erythromycin.

**Figure S2.**
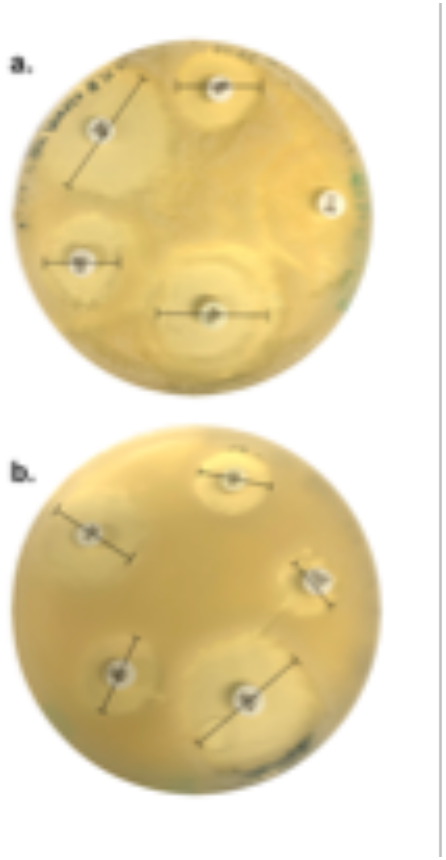
Selection of disk diffusion assay. **a.** Bt-kurstaki-1 with clindamycin, doxycycline, linezolid, sulfonamide/trimoxazole, and vancomycin. **b.** Bt-kurstaki-1 with clindamycin, doxycycline, linezolid, sulfonamide/trimoxazole, and vancomycin. Black bars indicate where measurement was taken.

**Supplemental Figure 3.**
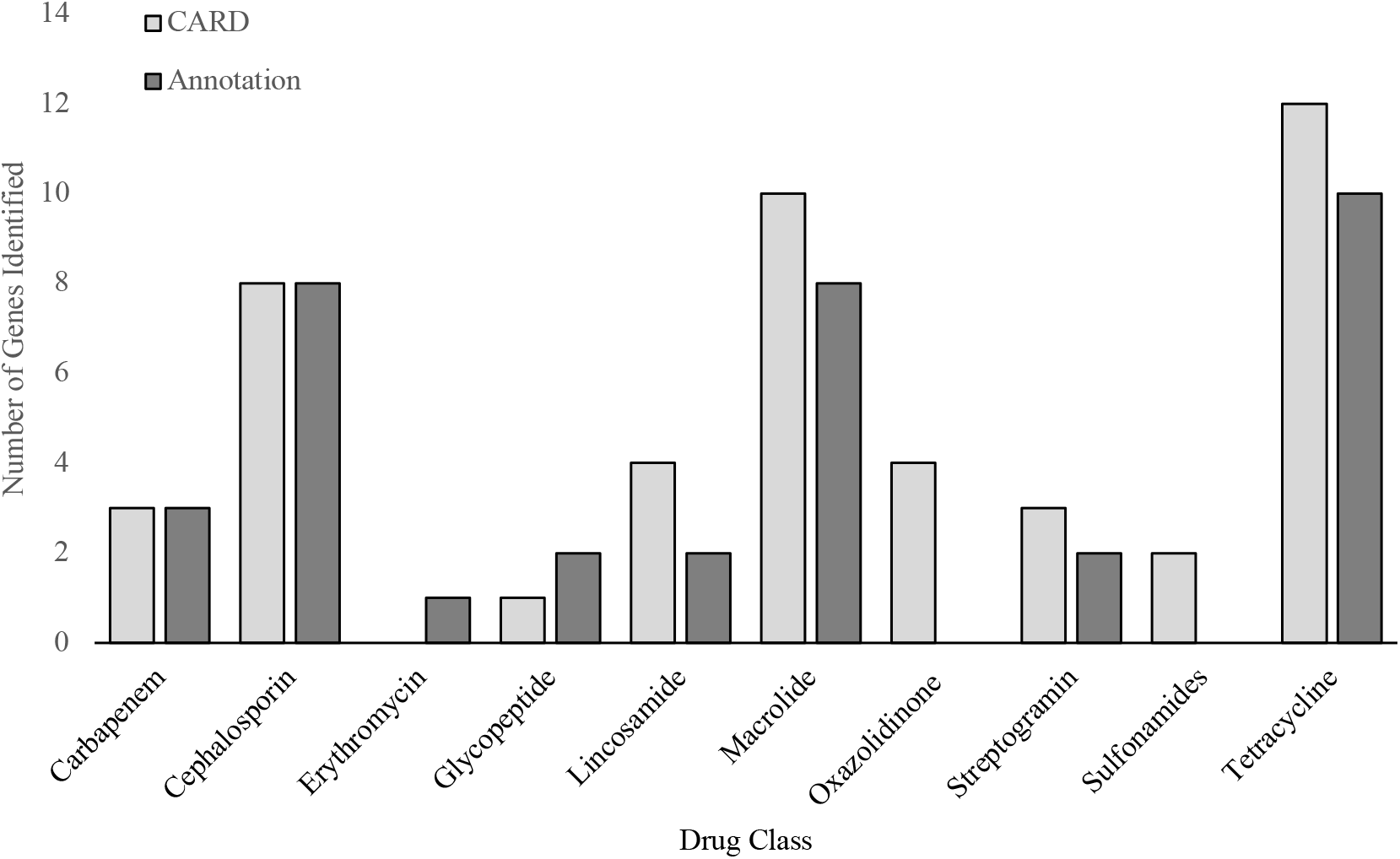
Comparison of Methods for Molecular Characterization by Drug Class Comparison of two methods, annotation review and homologous gene matching using a curated database (CARD), to determine similarity for molecular characterization of resistance phenotype.

